# The *Arabidopsis thaliana* carboxylesterase AtCXE12 converts volatile (*Z*)-3-hexenyl acetate to (*Z*)-3-hexenol

**DOI:** 10.1101/2023.03.14.532512

**Authors:** Tristan M. Cofer, Matthias Erb, James H. Tumlinson

## Abstract

The green leaf volatiles (*Z*)-3-hexenal, (*Z*)-3-hexenol, and (*Z*)-3-hexenyl acetate are produced by nearly all plants in response to wounding and insect attack, can be transferred between plants, metabolized, and act as defense cues. If and how plant leaves convert exogenous (*Z*)-3-hexenyl acetate to (*Z*)-3-hexenol is unknown. We show that Arabidopsis leaves rapidly convert exogenous (*Z*)-3-hexenyl acetate to (*Z*)-3-hexenol. Inhibitor and fractionation experiments identified the carboxylesterases AtCXE5 and AtCXE12 as likely contributors to (*Z*)-3-hexenyl acetate esterase activity in Arabidopsis leaves. Heterologous expression of AtCXE5 and AtCXE12 revealed that both enzymes hydrolyze (*Z*)-3-hexenyl acetate to (Z)-3-hexenol *in vitro*, and assays using T-DNA insertion mutant plants showed that AtCXE12 significantly contributes to the conversion of (*Z*)-3-hexenyl acetate to (*Z*)-3-hexenol *in planta*. Lastly, we found that leaves from several other plant species possess (*Z*)-3-hexenyl acetate esterase activity, as well as homologs of AtCXE5 and AtCXE12 from Arabidopsis. Collectively, our study provides a better understanding of green leaf volatile biosynthesis and conversion dynamics, necessary for unraveling the potential functions of these compounds.

## Introduction

In response to wounding and insect attack, nearly all plants produce green leaf volatiles (GLVs), including (*Z*)-3-hexenal, (*Z*)-3-hexenol, and (*Z*)-3-hexenyl acetate (Ameye et al., 2018; Matsui, 2006; Matsui & Engelberth, 2022). These compounds can act as a direct defense, with toxic or repellent effects on insect herbivores (Allmann et al., 2013; Hildebrand et al., 1993; Vancanneyt et al., 2001), and as an indirect defense, which attract natural enemies of an attacking herbivore (Allmann & Baldwin, 2010; Halitschke et al., 2008; Shiojiri et al., 2006). GLVs can also function as within- and between-plant signaling molecules, and enhance defenses in undamaged leaves (Ameye et al., 2015; Bate & Rothstein, 1998; Engelberth et al., 2004; Frost et al., 2008; Hu et al., 2019).

Upon wounding, plants rapidly produce large amounts of (*Z*)-3-hexenal (D’Auria et al., 2007). (*Z*)-3-Hexenal can react with amino and thiol groups in proteins (Anantharamkrishnan et al., 2020) and can thereby interfere with basic cellular processes. (*Z*)-3-Hexenal can be isomerized to (*E*)-2-hexenal by plants and some insect herbivores (Jones et al., 2022; Lin et al., 2022; Spyropoulou et al., 2017), resulting in the formation of isomers with distinct ecological functions (Allmann et al., 2013; Allmann & Baldwin, 2010). *In planta*, most of the (*Z*)-3-hexenal is converted to the less reactive (*Z*)-3-hexenol (Matsui et al., 2012). (*Z*)-3-Hexenol can then be glycosylated, to produce various (*Z*)-3-hexenyl glycosides, including (*Z*)-3-hexenyl vicianoside, a weakly toxic compound for insect herbivores (Sugimoto et al., 2014). (*Z*)-3-Hexenol is also converted to (*Z*)-3-hexenyl acetate, which is less polar and more volatile than its precursor, allowing it to diffuse more easily. Similar to other GLVs, (*Z*)-3-hexenyl acetate can be absorbed by nearby leaves to trigger defenses (Engelberth et al., 2004; Frost et al., 2008; Hu et al., 2019). Exogenous (*Z*)-3-hexenyl acetate is converted to (*Z*)-3-hexenyl glycosides, presumably via (*Z*)-3-hexenol, in receiving tissues (Ameye et al., 2020; Sugimoto et al., 2021). Understanding GLV biosynthesis and conversion dynamics is crucial to unravel the potential functions of these compounds.

GLV biosynthesis begins in the chloroplast of damaged cells, where lipoxygenases add molecular oxygen to the C_13_ position of linolenic acid (Christensen et al., 2013; Mochizuki et al., 2016). The resulting hydroperoxy fatty acid is then cleaved by hydroperoxide lyase to produce (*Z*)-3-hexenal (Matsui et al., 1996), which is reduced by an NADP(H)-dependent reductase in undamaged tissues to form (*Z*)-3-hexenol (Tanaka et al., 2018). (*Z*)-3-Hexenol can be further converted to (*Z*)-3-hexenyl acetate by an acetyl-CoA:(*Z*)-3-hexenol acetyltransferase (D’Auria et al., 2007) or glycosylated by UDP-dependent glycosyltransferases (Jing et al., 2019; Ohgami et al., 2015). Although (*Z*)-3-hexenyl acetate is often considered to be a stable end product, recent studies have shown that plants exposed to (*Z*)-3-hexenyl acetate accumulate (*Z*)-3-hexenyl glycosides (Ameye et al., 2020; Sugimoto et al., 2021), suggesting that (*Z*)-3-hexenyl acetate is hydrolyzed to (*Z*)-3-hexenol before further processing. Carboxylesterases that hydrolyze (Z)-3-hexenyl acetate have been identified in the fruits of several domesticated plants (Cao et al., 2019; Goulet et al., 2012; Martínez-Rivas et al., 2022; Souleyre et al., 2011). However, whether (*Z*)-3-hexenyl acetate is converted to (*Z*)-3-hexenol in plant leaves is unknown, and potential underlying enzymes remain unidentified.

Here, we aimed at determining if and how (*Z*)-3-hexenyl acetate can be transformed to (*Z*)-3-hexenol in *Arabidopsis thaliana* leaves. To this end, we used the Columbia ecotype, which, somewhat atypically, does not produce endogenous GLVs due to a mutation in hydroperoxide lyase (Duan et al., 2005). We discovered that Arabidopsis leaves rapidly convert (*Z*)-3-hexenyl acetate to (*Z*)-3-hexenol. We thus performed a series of inhibitor and fractionation experiments to identify esterases that catalyze this conversion. Finally, we used esterase mutant to confirm the involvement of the carboxylesterase AtCXE12 in (*Z*)-3-hexenol formation. We also determined that leaves from several other plant species possess (*Z*)-3-hexenyl acetate esterase activity alongside homologs of the active Arabidopsis carboxylesterases.

## Results

### Arabidopsis plants absorb and hydrolyze exogenous (*Z*)-3-hexenyl acetate

Given that plants exposed to (*Z*)-3-hexenyl acetate accumulate (*Z*)-3-hexenyl glycosides (Ameye et al., 2020; Sugimoto et al., 2021), we hypothesized that exogenous (*Z*)-3-hexenyl acetate is absorbed by plants and hydrolyzed to (*Z*)-3-hexenol. To test our hypothesis, we exposed Arabidopsis plants to synthetic (*Z*)-3-hexenyl acetate in an enclosed glass container, and measured the levels of (*Z*)-3-hexenyl acetate and (*Z*)-3-hexenol in leaves at different time points after exposure. We found that (*Z*)-3-hexenyl acetate and (*Z*)-3-hexenol accumulated in leaves within 5 min after (*Z*)-3-hexenyl acetate exposure (Figure 1). (*Z*)-3-Hexenyl acetate levels decreased between 5 and 30 min after exposure, and again between 30 and 180 min. (*Z*)-3-Hexenol levels increased between 5 and 10 min after exposure, and remained stable until decreasing between 30 and 180 min. Neither (*Z*)-3-hexenyl acetate nor (*Z*)-3-hexenol were detected in leaves after 90 min of exposure. These results show that Arabidopsis plants readily absorb and hydrolyze exogenous (*Z*)-3-hexenyl acetate to (*Z*)-3-hexenol as an intermediate compound, indicating the existence of a previously unidentified (*Z*)-3-hexenyl acetate esterase.

**Figure 1.**
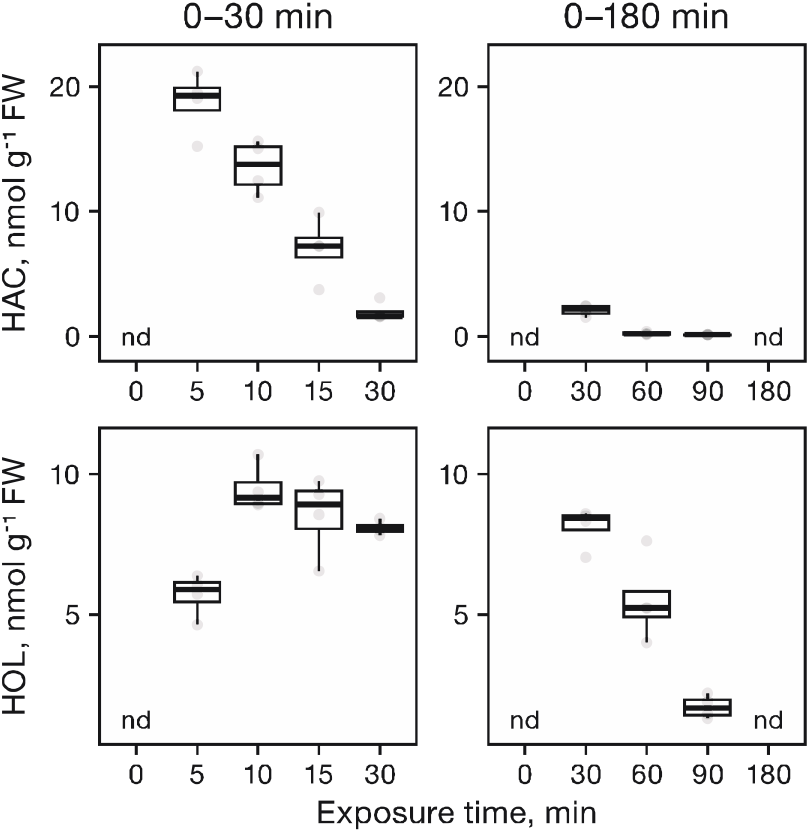
Levels of (*Z*)-3-hexenyl acetate and (*Z*)-3-hexenol in leaves from Arabidopsis plants exposed to (*Z*)-3-hexenyl acetate. Plants were exposed to synthetic (Z)-3-hexenyl acetate for 0, 5, 10, or 30 min (left column); and 0, 30, 60, 90, or 180 min (right column). (*Z*)-3-Hexenyl acetate (HAC) and (*Z*)-3-hexenol (HOL) were extracted from leaves and analyzed by GC-MS (n = 4). nd, not detected.

### Carboxylesterases contribute to (*Z*)-3-hexenyl acetate hydrolysis in Arabidopsis leaves

Carboxylesterases (EC 3.1.1.1) are a class of serine esterases that are reported to hydrolyze endogenous volatile esters in the fruit of both tomato (*Solanum lycopersicum*; Goulet et al., 2012) and peach (*Prunus persica*; Cao et al., 2019). To determine whether carboxylesterases also contribute to the hydrolysis of (*Z*)-3-hexenyl acetate in Arabidopsis leaves, we assayed extracts from Arabidopsis leaves for (*Z*)-3-hexenyl acetate esterase activity in the presence of the generic serine esterase inhibitor phenylmethylsulfonyl fluoride (PMSF) and the specific carboxylesterase inhibitor bis(*p*-nitrophenyl) phosphate (BNPP). We found that PMSF and BNPP each reduced the (*Z*)-3-hexenyl acetate esterase activity of leaf extracts in a dose-dependent manner (Figure 2). Average (*Z*)-3-hexenyl acetate esterase activity was reduced by 79% in the presence of 1 mM PMSF, and by 63% in the presence of 1 mM BNPP. Thus, carboxylesterases are expected to be a major contributor to the (*Z*)-3-hexenyl acetate esterase activity.

**Figure 2.**
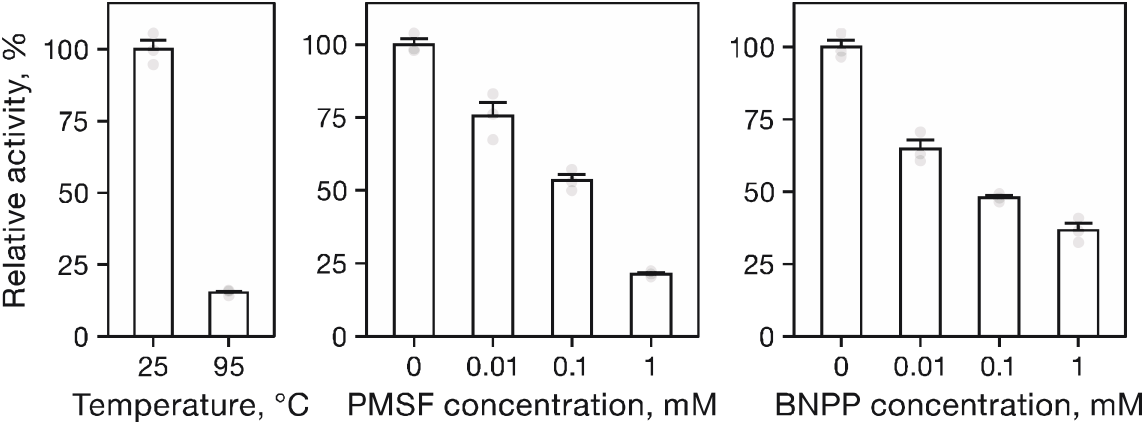
Relative (*Z*)-3-hexenyl acetate esterase activity of extracts from Arabidopsis leaves in the presence of the serine esterase inhibitor phenylmethylsulfonyl fluoride or the carboxylesterase inhibitor bis(*p*-nitrophenyl) phosphate. Leaf extracts were incubated with 0, 0.01, 0.1, or 1 mM phenylmethylsulfonyl fluoride (PMSF) or inhibitor bis(*p*-nitrophenyl) phosphate (BNPP) at 23 °C for 1 h, after which extracts were assayed for (*Z*)-3-hexenyl acetate esterase activity (n = 3). Activities are presented as a percentage relative to 0 mM PMSF/BNPP (100%). Data are mean ± SEM.

### AtCXE5 and AtCXE12 are (*Z*)-3-hexenyl acetate esterase candidates

The Arabidopsis genome contains 20 carboxylesterase genes, 18 of which are expressed in leaves (Marshall et al., 2003). To identify the specific carboxylesterases that contribute to (*Z*)-3-hexenyl acetate hydrolysis in Arabidopsis leaves, we purified the (*Z*)-3-hexenyl acetate esterase activity from leaf extracts using a three-step purification procedure (Table 1). Leaf extracts were first fractionated by ammonium sulfate precipitation, and the fraction precipitating between 40 and 80% ammonium sulfate saturation was purified on a phenyl Sepharose column. The elution profile from the phenyl Sepharose column revealed that the (*Z*)-3-hexenyl acetate esterase activity was distributed across three unequal pools (Figure 3). The largest pool, which contained approximately 80% of the total (*Z*)-3-hexenyl acetate esterase activity recovered at this step, was resolved on a Mono Q column to a single peak. Attempts to further purify the (*Z*)-3-hexenyl acetate esterase activity were unsuccessful, resulting in a substantial decrease in yield, with only modest gains in purity (data not shown). Thus, proteins contained in the three most active fractions collected from the Mono Q column were combined and subjected to proteomic analysis. A total of 85 Arabidopsis proteins with two or more peptides were identified, including two putative carboxylesterases: AtCXE5 and AtCXE12 (Supplemental Table S2; Supplemental Figure S1). Notably, both AtCXE5 and AtCXE12 were previously shown to hydrolyze other acetate esters *in vitro* (Gershater et al., 2007), and were therefore considered as strong candidates for further study.

**Table 1.** Purification of (*Z*)-3-hexenyl acetate esterase activity from Arabidopsis leaves.

| Purification step | Protein | Total activity | Specific activity | Purification | Yield |
| --- | --- | --- | --- | --- | --- |
|  | mg | pkat | pkat mg<sup>-1</sup> | fold | % |
| Crude extract | 364 | 1848 | 5.08 | 1.00 | 100 |
| Ammonium sulfate | 102 | 1079 | 10.5 | 2.08 | 58 |
| Phenyl Sepharose | 2.03 | 683 | 336 | 66.3 | 37 |
| Mono Q | 0.23 | 194 | 838 | 165 | 10 |

**Figure 3.**
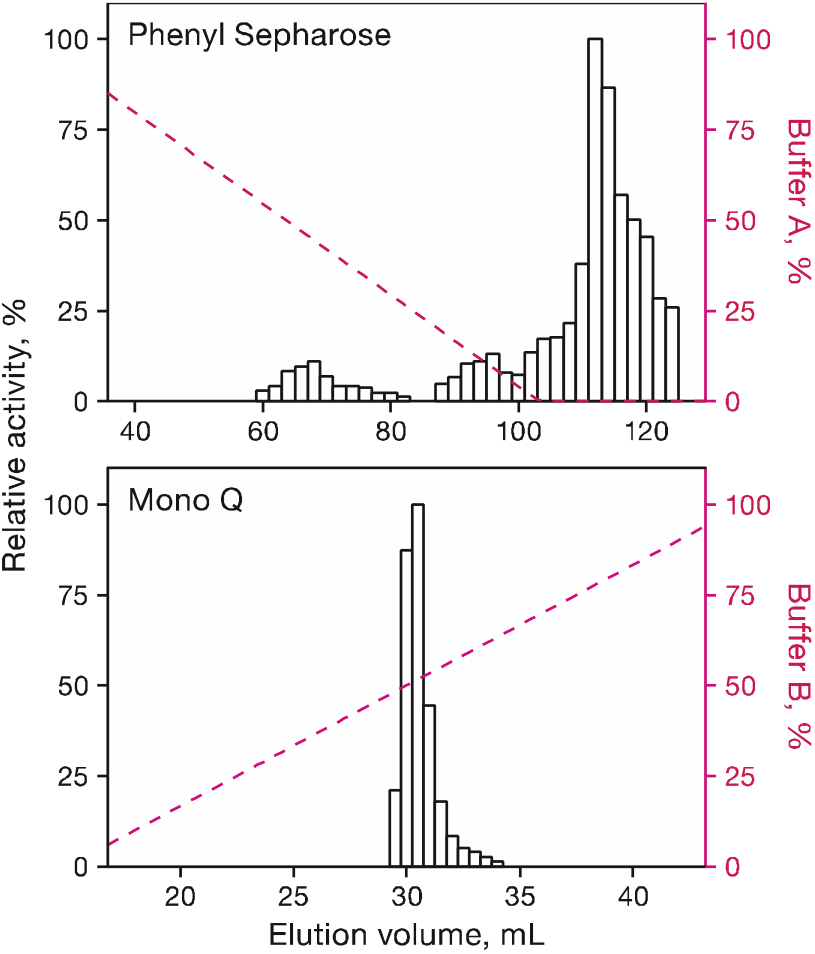
Purification of (Z)-3-hexenyl acetate esterase activity from Arabidopsis leaves. Elution profiles of (Z)-3-hexenyl acetate esterase activity from phenyl Sepharose and Mono Q columns.

### Recombinant AtCXE5 and AtCXE12 hydrolyze (*Z*)-3-hexenyl acetate

The full-length coding sequences of *AtCXE5* and *AtCXE12* were heterologously expressed in *Escherichia coli*, and the recombinant proteins were assayed for esterase activity with (*Z*)-3-hexenyl acetate and several other (Z)-3-hexenyl esters (Table 2). Both AtCXE5 and AtCXE12 exhibited relatively high activity with (*Z*)-3-hexenyl acetate, (*Z*)-3-hexenyl propionate, and (*Z*)-3-hexenyl butyrate; and little to no activity with (*Z*)-3-hexenyl isobutyrate, (*Z*)-3-hexenyl 2-methylbutyrate, and (*Z*)-3-hexenyl tiglate. Both recombinant proteins had an optimum pH of 7.5 in Tris-HCl buffer. Kinetic analyses revealed that AtCXE5 and AtCXE12 each preferred (*Z*)-3-hexenyl butyrate as substrate over (*Z*)-3-hexenyl propionate and (*Z*)-3-hexenyl acetate (Table 3). The apparent K_m_ value of AtCXE5 for (*Z*)-3-hexenyl acetate was 1.45-times higher than that of AtCXE12, whereas the k_cat_ values of AtCXE5 and AtCXE12 for (*Z*)-3-hexenyl acetate were similar. Overall, the catalytic efficiency (k_cat_/K_m_) of AtCXE5 for (*Z*)-3-hexenyl acetate was 1.68-times lower than that of AtCXE12. Thus, both AtCXE5 and AtCXE12 can exhibit (*Z*)-3-hexenyl acetate esterase activity *in vitro*.

**Table 2.** Substrate specificity of AtCXE5 and AtCXE12.

| Substrate | Relative activity, % <sup>†</sup> |  |
| --- | --- | --- |
|  | AtCXE5 | AtCXE12 |
| (Z)-3-hexenyl acetate | 100.00 ± 8.88 | 100.00 ± 14.17 |
| (Z)-3-hexenyl propionate | 57.59 ± 8.37 | 180.72 ± 15.54 |
| (Z)-3-hexenyl butyrate | 101.78 ± 4.65 | 299.33 ± 5.36 |
| (Z)-3-hexenyl isobutyrate | 5.29 ± 0.59 | 6.65 ± 1.13 |
| (Z)-3-hexenyl 2-methylbutyrate | nd | 6.21 ± 1.15 |
| (Z)-3-hexenyl tiglate | nd | nd |

<sup>†</sup> All assays were performed under identical reaction conditions. Activates are presented as a percentage relative to (Z)-3-hexenyl acetate (100%). Data are mean ± SEM (n = 3). nd, nondetectable activity (< 2%)

**Table 3.**
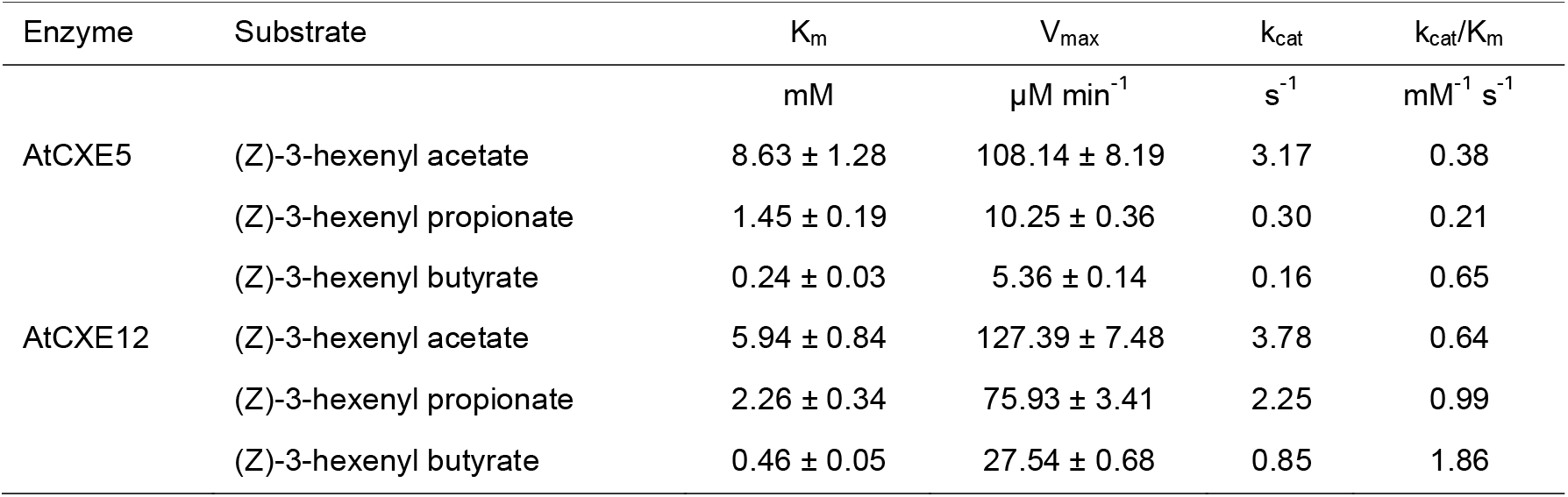
Kinetic parameters of AtCXE5 and AtCXE12.

| Enzyme | Substrate | K<sub>m</sub> | V<sub>max</sub> | k<sub>cat</sub> | k<sub>cat</sub>/K<sub>m</sub> |
| --- | --- | --- | --- | --- | --- |
|  |  | mM | μM min<sup>-1</sup> | s<sup>-1</sup> | mM<sup>-1</sup> s<sup>-1</sup> |
| AtCXE5 | (Z)-3-hexenyl acetate | 8.63 ± 1.28 | 108.14 ± 8.19 | 3.17 | 0.38 |
|  | (Z)-3-hexenyl propionate | 1.45 ± 0.19 | 10.25 ± 0.36 | 0.30 | 0.21 |
|  | (Z)-3-hexenyl butyrate | 0.24 ± 0.03 | 5.36 ± 0.14 | 0.16 | 0.65 |
| AtCXE12 | (Z)-3-hexenyl acetate | 5.94 ± 0.84 | 127.39 ± 7.48 | 3.78 | 0.64 |
|  | (Z)-3-hexenyl propionate | 2.26 ± 0.34 | 75.93 ± 3.41 | 2.25 | 0.99 |
|  | (Z)-3-hexenyl butyrate | 0.46 ± 0.05 | 27.54 ± 0.68 | 0.85 | 1.86 |

### AtCXE12 contributes to the hydrolysis of exogenous (*Z*)-3-hexenyl acetate *in planta*

To investigate whether AtCXE5 and AtCXE12 contribute to the hydrolysis of (*Z*)-3-hexenyl acetate *in planta*, we searched the T-DNA Express database (http://signal.salk.edu/cgi-bin/tdnaexpress) for T-DNA insertion alleles in each locus. We identified two *atcxe5* T-DNA insertion alleles, each with the T-DNA insertion located in the 5’UTR (Supplemental Figure S2). However, RT-PCR analysis revealed no difference in the levels of *AtCXE5* transcripts between T-DNA insertion mutants and the WT. Thus, these lines were not considered for further experiments. For AtCXE12, a single null allele was identified, with the T-DNA located in the coding region, 102 bp downstream from the start codon (Supplemental Figure S2). This line was thus used for further experiments.

Leaf extracts from the *atcxe12* T-DNA insertion mutant exhibited a 52% reduction in average (*Z*)-3-hexenyl acetate esterase activity relative to those from the WT (Figure 4). Given that the specific carboxylesterase inhibitor BNPP reduces the average (*Z*)-3-hexenyl acetate esterase activity of leaf extracts by approximately 63%, we reasoned that AtCXE12 should be the major esterase that converts (*Z*)-3-hexenyl acetate to (*Z*)-3-hexenol. To quantify residual esterase activity in the *atcxe12* null mutant, we measured the reduction of (*Z*)-3-hexenyl acetate conversion in the presence of 1 mM BNPP. We found that BNPP reduced the average (*Z*)-3-hexenyl acetate esterase activity of *atcxe12* leaf extracts by 25% only, thus indicating that AtCXE12 is indeed the dominant (*Z*)-3-hexenyl acetate esterase in Arabidopsis (Figure 4).

**Figure 4.**
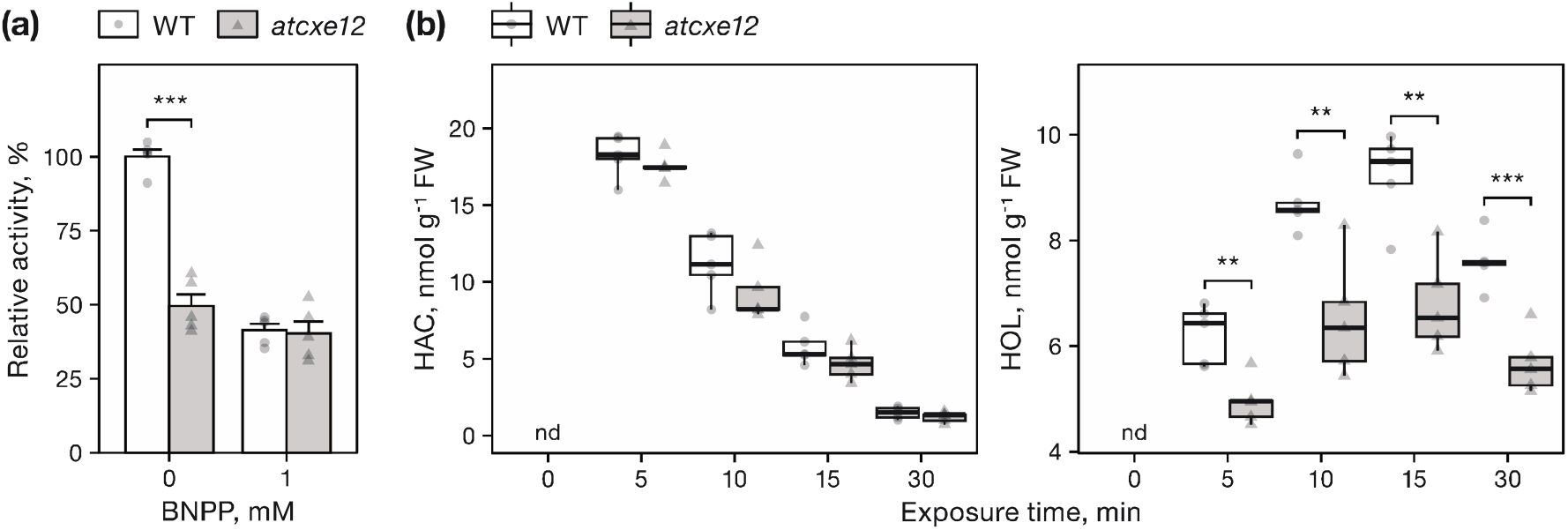
Comparison of atcxe12 and WT plants. **(a)** Relative (*Z*)-3-hexenyl acetate esterase activity of extracts from *atcxe12* and WT leaves. Leaf extracts were incubated with 0 or 1 mM bis(*p*-nitrophenyl) phosphate (BNPP) at 23 °C for 1 h, after which extracts were assayed for (*Z*)-3-hexenyl acetate esterase activity. Activities are presented as a percentage relative to the WT with 0 mM BNPP (100%). Data are mean ± SEM (n = 5). **(b)** Levels of (*Z*)-3-hexenyl acetate (HAC) and (*Z*)-3-hexenol (HOL) in leaves from Arabidopsis plants exposed to (*Z*)-3-hexenyl acetate. Plants were exposed to synthetic (*Z*)-3-hexenyl acetate for 0, 5, 10, or 30 min. (*Z*)-3-Hexenyl acetate and (*Z*)-3-hexenol were extracted from leaves and analyzed by GC-MS. Data are mean ± SEM (n = 5). Asterisks represent significant differences from WT plants (\*\**P* < 0.01; \*\*\**P* < 0.001; Students *t*-test). nd, not detected.

To further investigate the contribution of AtCXE12 to (*Z*)-3-hexenyl acetate hydrolysis in intact Arabidopsis leaves, we measured the levels of (*Z*)-3-hexenyl acetate and (*Z*)-3-hexenol in leaves from *atcxe12* and WT plants following exposure to synthetic (*Z*)-3-hexenyl acetate. We found that (*Z*)-3-hexenol levels were consistently lower in leaves from *atcxe12* plants than in those from WT plants after (*Z*)-3-hexenyl acetate exposure (Figure 4). The difference in average (*Z*)-3-hexenol levels between *atcxe12* and WT plants was approximately 25% across time points.

### CXE5 and CXE12 are conserved across different plant species

Lastly, we investigated whether AtCXE5 and AtCXE12 are conserved across different plant species and whether their presence is associated with (*Z*)-3-hexenyl acetate esterase activity. We found that leaf extracts from common bean (*Phaseolus vulgaris*), tomato, wheat (*Triticum aestivum*), and maize (*Zea mays*) were able to convert (*Z*)-3-hexenyl acetate to (*Z*)-3-hexenol (Figure 5). Homologs sharing greater than 40% amino acid sequence identity with AtCXE5 and AtCXE12 were also identified in each species (Figure 5), suggesting that these enzymes are conserved across the plant kingdom similarly to the capacity of plants to hydrolyze (*Z*)-3-hexenyl acetate.

**Figure 5.**
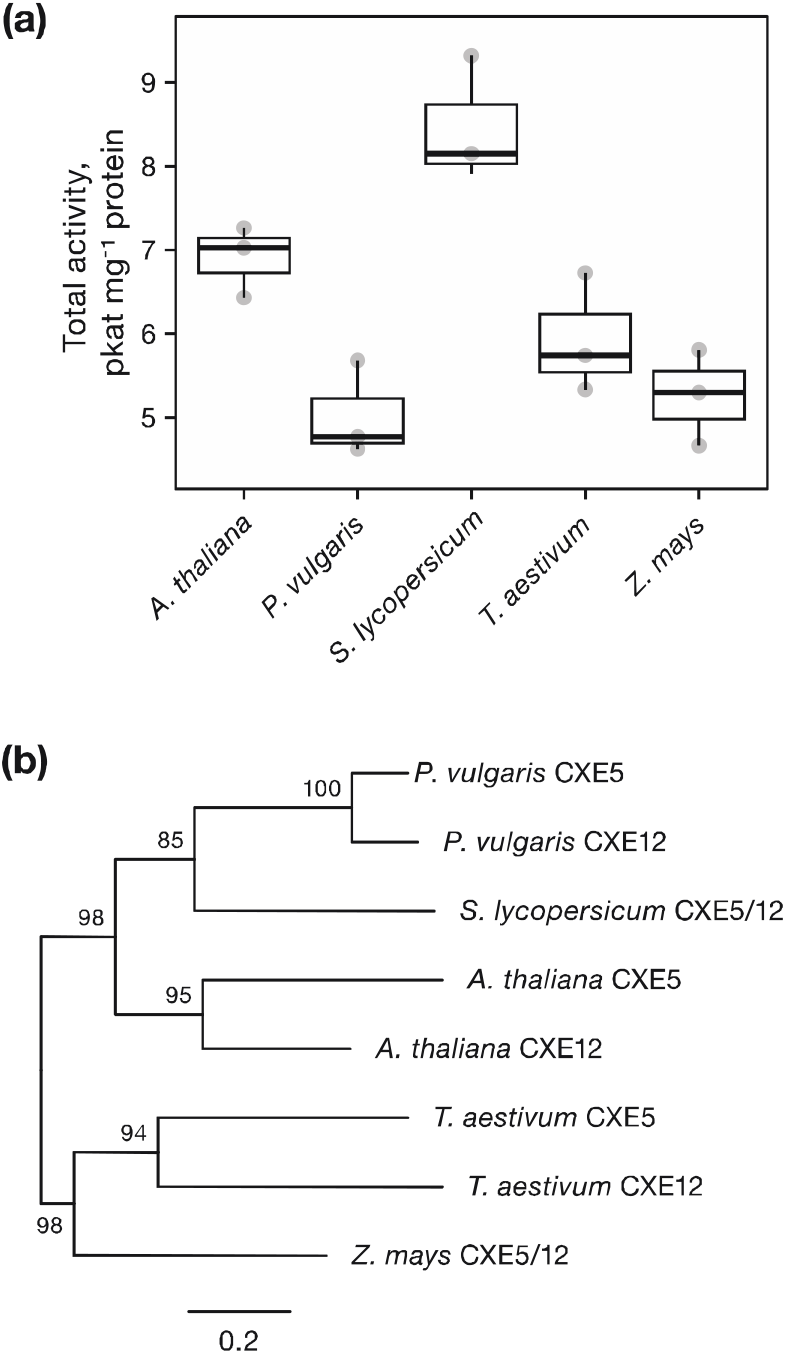
(*Z*)-3-Hexenyl acetate esterase activity and identification of AtCXE5 and AtCXE12 homologs in different plant species. **(a)** (*Z*)-3-Hexenyl acetate esterase activity of leaf extracts from *Arabidopsis thaliana, Phaseolus vulgaris, Solanum lycopersicum, Triticum aestivum, Zea mays*. Data are mean ± SEM (n = 3). **(b)** Phylogenetic analysis of AtCXE5 and AtCXE12 homologs. Amino acid sequences with the highest similarity to AtCXE5 and AtCXE12 from Arabidopsis were obtained using BLASTP on the Phytozome website. Sequences were aligned using MAFFT v7.401, and a maximum likelihood tree was constructed using IQ-TREE v2.0.3 with 1,000 ultrafast bootstrap replicates. The scale bare represents the number of amino acid substitutions per site.

## Discussion

(*Z*)-3-Hexenyl acetate is commonly converted to (*Z*)-3-hexenyl glycosides in plant leaves, which may have important consequences for its biological functions and effects. However, if and how this conversion happens via (*Z*)-3-hexenol formation, remained unknown. Here, we show that (*Z*)-3-hexenyl acetate is rapidly converted to (*Z*)-3-hexenol in plant leaves, and identify two carboxylesterases that catalyze this reaction.

Carboxylesterases have been found to hydrolyze volatile esters in fruit of several domesticated plants. SlCXE1 from tomato for instance, is preferentially expressed in fruit and can hydrolyze (*Z*)-3-hexenyl acetate and other volatile esters that contribute to fruit flavor (Goulet et al., 2012). MdCXE1 from apple (*Malus* × *domestica*; Souleyre et al., 2011) PpCXE1 from peach (Cao et al., 2019), and FanCXE1 from strawberry (*Fragaria* × *ananassa*; Martínez-Rivas et al., 2022) are similarly associated with the hydrolysis of volatile esters during fruit ripening. Here, we show that carboxylesterases also contribute to the (*Z*)-3-hexenyl acetate esterase activity in Arabidopsis leaves. We found that AtCXE5 and AtCXE12 hydrolyze (*Z*)-3-hexenyl acetate and exhibit high relative activity with longer, straight chain (Z)-3-hexenyl esters. The overall kinetic parameters of AtCXE5 and AtCXE12 were similar to those reported for other carboxylesterases (http://brenda-enzymes.info). However, the apparent K_m_ values of AtCXE5 and AtCXE12 for (*Z*)-3-hexenyl acetate were both higher than those of SlCXE1 (K_m_ = 0.47) and PpCXE1 (K_m_ = 0.99). These differences could be accounted for by several factors. For example, AtCXE5 and AtCXE12 may be adapted to function efficiently at relatively high (*Z*)-3-hexenyl acetate concentrations, such as those found near damaged leaves (D’Auria et al., 2007). Consumer preferences during domestication may also have led to selection for alleles that result in lower (*Z*)-3-hexenyl acetate content in ripening fruits. As a result, carboxylesterases in fruit of some domesticated plants could have evolved a stronger affinity for (*Z*)-3-hexenyl acetate and other volatile esters that are disliked by consumers.

While some carboxylesterases exhibit activity with only a few substrates, others are able to hydrolyze a broad range of esters (Marshall et al., 2003). This is the case with AtCXE5 and AtCXE12, which hydrolyze (*Z*)-3-hexenyl acetate, as well as esters that are not known to be produced by Arabidopsis, such as (*Z*)-3-hexenyl butyrate and (*Z*)-3-hexenyl propionate. Both enzymes were also reported to hydrolyze the model acetate ester *p*-nitrophenyl acetate, and AtCXE12 alone was shown to convert methyl-2,4-dichlorophenoxyacetate to the active herbicide 2,4-dichlorophenoxyacetic acid (Gershater et al., 2007). These findings suggest that AtCXE5 and AtCXE12 may hydrolyze a number of different esters in Arabidopsis leaves.

What is the biological significance of converting exogenous (*Z*)-3-hexenyl acetate to (*Z*)-3-hexenol in plant leaves? One possibility is that hydrolyzing (*Z*)-3-hexenyl acetate may prevent it from binding to an as yet unidentified receptor, thereby buffering (*Z*)-3-hexenyl acetate-induced defense responses. Another possibility is that hydrolysis serves to activate (*Z*)-3-hexenyl acetate by converting it to (*Z*)-3-hexenol, which then induces plant defenses. This second possibility is similar to the process by which plants hydrolyze volatile methyl salicylate and methyl jasmonate to the phytohormones salicylic acid and jasmonic acid, respectively (Forouhar et al., 2005; Koo et al., 2013). In both cases, silencing the specific methylesterase responsible for hydrolyzing methyl salicylate and methyl jasmonate attenuated plant responses to these volatile signaling molecules (Manosalva et al., 2010; Park et al., 2007; Wu et al., 2008). We found that leaves from *atcxe12* T-DNA insertion mutants accumulate significantly lower levels of (*Z*)-3-hexenol compared to those from WT plants following exposure to (*Z*)-3-hexenyl acetate. However, the approximately 25% difference in average (*Z*)-3-hexenol levels observed between *atcxe12* and WT plants indicates that other (*Z*)-3-hexenyl acetate esterases remain to be discovered in Arabidopsis. Further work, including the creation of higher-order mutants and overexpression lines will help to generate the necessary molecular tools to explore the biological significance of (*Z*)-3-hexenol hydrolysis. Transferring the knowledge gained here to plants that show strong robust defense response to (Z)-3-hexenyl acetate will be essential in this context.

## Materials and methods

### Plant material and growth conditions

The Arabidopsis (*Arabidopsis thaliana* (L.) Heynh.) ecotype Columbia (Col-0) was used as the wild type (WT) for all experiments. The T-DNA insertion lines *atcxe5-1* (SALK_209596C), *atcxe5-2* (SALK_106155C1), and *atcxe12* (SAIL_445_G03) were obtained from the Arabidopsis Biological Resource Center (Ohio State University, Columbus, OH, USA). All T-DNA insertion lines were genotyped using a T-DNA-specific primer and gene-specific primer pairs flanking each insertion site (Supplemental Table S1).

Plants were grown on a mixture of PRO-MIX® FLX (Premier Horticulture Inc., Quakertown, PA, USA) and perlite (3:1, v/v) supplemented with Osmocote® Smart Release® 14-14-14 plant food (Scotts Company LLC, Marysville, OH, USA) in a growth chamber (22 °C) under a short-day photoperiod (10:14, light/dark) with illumination from cool-white fluorescent lamps (∼120 μmol photons m^-2^ s^-1^). Four- to five-week-old plants in the vegetative stage were used for all experiments.

### (*Z*)-3-hexenyl acetate exposure treatments

To expose Arabidopsis plants to (*Z*)-3-hexenyl acetate, we enclosed individual plants in a 500-cm^3^ glass container along with a cotton swab on which we applied 0.5 μmol of synthetic (*Z*)-3-hexenyl acetate dissolved in 5 μL of dichloromethane. Whole rosettes were harvested at different time points after exposure and flash-frozen in liquid nitrogen. Samples were ground in liquid nitrogen and stored at −80 °C until use.

### GLV quantification

GLVs were extracted from Arabidopsis leaves following a protocol modified from Seidl-Adams et al. (2015). Briefly, 200 mg of ground plant material were homogenized in 0.8 mL of n-hexane using a FastPrep F120 tissue homogenizer (Qbiogene, Carlsbad, CA, USA). The homogenate was centrifuged at 15,000 *g* for 3 min, and the upper organic phase was transferred to a 4-mL glass vial. Volatiles were collected onto a SuperQ filter trap (20 mg; Alltech Associates, Deerfield, IL, USA) by vapor-phase extraction as described in Schmelz et al. (2004). The trapped volatiles were eluted with 100 μL of n-hexane/dichloromethane (1:1, v/v) containing 500 ng of nonyl acetate as an internal standard. Samples were analyzed on an Agilent 6890 series gas chromatograph (GC) coupled to an Agilent 5973 quadrupole mass selective detector (MS; interface temperature, 250 °C; quadrupole temperature, 150 °C; source temperature, 230 °C; electron energy, 70 eV). An aliquot of each sample was injected in splitless mode onto an HP-5MS column (30 m × 0.25 mm i.d. × 0.25 μm thickness; Agilent, Palo Alto, CA, USA). Helium was used as the carrier gas at a constant flow rate of 1 mL min^-1^. The oven temperature was held at 40 °C for 2 min, then increased to 280 °C at 15 °C min^-1^ and held at 280 °C for 3 min. Mass spectra were acquired in scan mode from 50 to 300 m/z. Samples were normalized by dividing the area of the base peak for each analyte by that of the internal standard. Absolute quantities were obtained using a standard curve.

### Protein extraction and esterase activity assays

Ground plant material was homogenized in 10 volumes (w/v) of ice-cold extraction buffer (50 mM Tris-HCl, pH 7.5, 2 mM dithiothreitol) using a micro-pestle. The homogenate was centrifuged at 15,000 *g* for 10 min, and the supernatant was retained for esterase activity assays. Protein concentration was determined using the Bradford method (Bradford, 1976) with bovine serum albumin as a standard.

Esterase activity assays were carried out in a 200-μL reaction mixture containing 50 mM of Tris-HCl buffer (pH 7.5), 0.2 mM of volatile ester substrate, and varying amounts of protein solution. Reaction mixtures were incubated at 30 °C for 20 min, after which the products formed were extracted and analyzed by GC-MS. To measure the effects of the generic serine esterase inhibitor phenylmethylsulfonyl fluoride (PMSF) and the specific carboxylesterase inhibitor bis(*p*-nitrophenyl) phosphate (BNPP) on (*Z*)-3-hexenyl acetate esterase activity, we incubated extracts from Arabidopsis leaves with varying concentrations of each inhibitor at 23 °C for 1 h, and assayed the inhibitor-treated extracts for (*Z*)-3-hexenyl acetate esterase activity under standard assay conditions.

### (*Z*)-3-hexenyl acetate esterase purification and identification

Approximately 100 g of ground plant material was homogenized in 150 mL of ice-cold extraction buffer (50 mM Tris-HCl, pH 7.5, 2 mM dithiothreitol) using a Waring Blendor. The homogenate was filtered through three layers of muslin to remove debris, and the filtrate was centrifuged at 10,000 *g* for 20 min to afford a crude extract as the supernatant. The crude extract was fractionated with ammonium sulfate between 40 and 80% saturation, and the precipitate was dissolved in buffer A (50 mM Tris-HCl, pH 7.5, 0.5 M ammonium sulfate). The protein solution was applied to a 5-mL phenyl Sepharose HP column (Cytiva, Marlborough, MA, USA) equilibrated with buffer A. The column was washed with 25 mL of buffer A, and bound proteins were eluted over an 80-mL decreasing linear gradient to 0 mM ammonium sulfate at a flow rate of 1 mL min^-1^. Fractions were collected at 2-mL intervals and assayed for (*Z*)-3-hexenyl acetate esterase activity. Active fractions were combined and exchanged into buffer B (50 mM Tris-HCl, pH 7.5) before being applied to a Mono Q™ HR 5/5 column (GE Healthcare, Piscataway, NJ, USA) equilibrated with the same buffer. The column was washed with 15 mL of buffer B, and bound proteins were eluted over a 30-mL increasing linear gradient to 0.2 M NaCl at a flow rate of 0.5 mL min^-1^. Fractions were collected at 0.5-mL intervals and assayed for (*Z*)-3-hexenyl acetate esterase activity. Active fractions were combined and subjected to proteomic analysis by liquid chromatography tandem mass spectrometry (LC-MS/MS).

For proteomic analysis, proteins were alkylated with iodoacetamide and digested with Trypsin/Lys-C Mix (Promega, Madison, WI, USA), according to the manufacturer’s instructions. Digested peptides were analyzed on a Bruker nanoElute® liquid chromatography system coupled to a Bruker timsTOF fleX™ mass spectrometer operated in PASEF mode with a capillary voltage of 1.6 kV. Peptides were separated on a reversed-phase C_18_ nano column (15 cm × 0.75 μm i.d., 1.7 μm particle size, 100 Å pore size; PepSep, Marslev, DK) maintained at 50 °C. Water/formic acid (100:0.1, v/v) and acetonitrile/formic acid (100:0.1, v/v) were employed as mobile phases A and B, respectively. The elution profile was as follows: 0–17.22 min, 5%–30% A in B; 17.22–17.72 min, 30%–95% A in B; 17.72–20.85, 95% A in B; and 20.85–30 min 5% A in B, with a flow rate of 0.3 μL min^-1^. Mass spectra were acquired from 100 to 1,700 m/z. Data were analyzed against the NCBI RefSeq *Arabidopsis thaliana* database plus a list of 536 common laboratory contaminants using Byonic™ software (v3.6.0; Protein Metrics Inc, Cupertino, CA, USA).

### RNA extraction and cDNA synthesis

Total RNA was extracted from Arabidopsis leaves using the RNeasy Mini Plant Kit (Qiagen, Valencia, CA, USA) and treated with TURBO™ DNase using the TURBO DNA-free™ Kit (Applied Biosystems, Carlsbad, CA, USA), each according to the manufacturer’s instructions. First-strand cDNA was synthesized from 1–2 μg of DNase-treated total RNA using SuperScript™ IV Reverse Transcriptase (Invitrogen, Carlsbad, CA, USA) and a mixture of anchored oligo(dT)_18_ and random octamer primers (Genomics Core Facility, The Pennsylvania State University, University Park, PA, USA).

### Cloning and heterologous expression

The full-length coding sequences of *AtCXE5* and *AtCXE12* were amplified from Arabidopsis cDNA using gene-specific primers (Supplemental Table S1) and ligated into the pET28b expression vector (Novagen, Madison, WI, USA) in-frame with a N-terminal hexahistidine (His)-tag. Individual expression constructs were subsequently transformed into *Escherichia coli* Rosetta™ 2 (DE3)pLysS cells (Novagen) for heterologous expression. Cells harboring an expression construct were grown at 37 °C in Lysogeny Broth supplemented with kanamycin (50 μg mL^-1^) and chloramphenicol (50 μg mL^-1^) to an OD_600_ of 0.7–0.8. Cultures were cooled to 23 °C, and β-D-1-thiogalactopyranoside was added to a final concentration of 1 mM. Induced cultures were incubated at 16 °C with continuous shaking (200 rpm) for 20 h, after which cells were harvested by centrifugation and stored at −80 °C until use.

### Recombinant protein purification

Frozen cell pellets were thawed on ice and resuspended in lysis buffer (50 mM Tris-HCl, pH 7.5, 500 mM NaCl, ∼10 μg mL^-1^ DNase I). The suspension was sonicated five times for 30-s intervals at 40 kHz, and the lysate was centrifuged at 100,000 *g* for 30 min. The supernatant was applied to a TALON® metal affinity resin column (Takara Bio, Mountain View, CA, USA), and the column was washed with 10 column volumes of wash buffer (50 mM Tris-HCl, pH 7.5, 500 mM NaCl, 5 mM imidazole, 2 mM β-mercaptoethanol, 5 mM MgCl_2_, 5 mM ATP). Proteins were eluted with a minimal volume of elution buffer (50 mM Tris-HCl, pH 7.5, 500 mM NaCl, 150 mM imidazole, 5 mM β-mercaptoethanol) and exchanged into storage buffer (50 mM Tris-HCl, pH 7.5, 150 mM NaCl, 10% (w/v) glycerol, 2 mM dithiothreitol).

### Enzyme kinetics

Protein concentration and reaction conditions were chosen such that the reaction velocity was linear during the incubation period. The pH optimum was determined in 50 mM of citrate buffer (pH 4.0−6.0), 50 mM of sodium phosphate buffer (pH 6.0−7.5), or 50 mM of Tris-HCl buffer (pH 7.5−9.0). Standard assays were carried out in a 200-μL reaction mixture containing 50 mM of Tris-HCl buffer (pH 7.5), 5– 16,000 μM of substrate, and 0.5 μM (∼4 μg) of affinity-purified recombinant protein. Reaction mixtures were incubated at 30 °C for 10 min, after which the products formed were extracted and analyzed by GC-MS. All reactions were assayed in triplicate. Data were fitted to the Michaelis-Menten equation by non-linear regression.

### Phylogenetic analysis

The Phytozome BLASTP tool was used to identify the closest homologs of AtCXE5 and AtCXE12 from *Phaseolus vulgaris* (v2.1), *Solanum lycopersicum* (ITAG4.0), *Triticum aestivum* (v2.2), and *Zea mays* (RefGen_V4). Selected amino acid sequences were aligned using MAFFT v7.401 (default settings; Katoh & Standley, 2013). ModelFinder was used to obtain the best-fit substitution model based on BIC values (Kalyaanamoorthy et al., 2017), and a maximum likelihood tree was constructed using IQ-TREE v2.0.3 (Minh et al., 2020) with 1,000 ultrafast bootstrap replicates (UFBoot2; Hoang et al., 2018).

## Acknowledgements

The authors would like to extend their appreciation to Anne Stanley for her expertise in proteomic analysis, and to Irmgard Seidl-Adams, Tim Moural, and Rose Fang for their instrumental role in the completion of this research. We thank Jurgen Engelberth, Gary Felton, and John Tooker for their valuable feedback on an earlier version of this manuscript. This paper is dedicated to James H. “Jim” Tumlinson, who passed away during the final preparations of this manuscript.

