## Supplemental materials for "The *Arabidopsis thaliana* carboxylesterase AtCXE12 converts volatile (*Z*)-3-hexenyl acetate to (*Z*)-3-hexenol"

AtCXE5 (At1g49660)

MES**EIASEFL PFCRIYKDGR VERLIGTDTI PASLDPTYDV VSKDVIYSPE NNLSVRLFLP** 60

**HKSTK**LTAGN KLPLLIYIHG GAWIIESPFS PLYHNYLTEV VKSANCLAVS VQYRRAPEDP 120

VPAAYEDVWS AIQWIFAHSN GSGPVDWINK HADFG**KVFLG GDSAGGNISH HMAMKA**GKEK 180

KLDLKIKGIA VVHPA**FWGTD PVDEYDVQDK ETRS**GIAEIW EKIASPNSVN GTDDPLFNVN 240

GSGSDFSGLG CDKV**LVAVAG KDVFVRQGLA YAAKLEK**CEW EGTV**EVVEEE GEDHVFHLQN** 300

**PKSDKA**LKFL KKFVEFIIG 319

AtCXE12 (At3g48690)

MDS**EIAVDCS PLLKIYKSGR IERLMGEATV PPSSEPQNGV VSKDVVYSAD NNLSVRI**YLP 60

EKAAAETDSK LPLLVYFHGG GFIIETAFSP TYHTFLTTSV SASNCVAVSV DY**RRAPEHPI** 120

**SVPFDDSW**TA LKWVFTHITG SGQEDWLNKH ADFS**RVFLSG DSAGANIVHH MAMRA**AKEKL 180

SPGLNDTGIS GII**LLHPYFW SKTPIDEKDT KDETLRM**KIE AFWMMASPNS KDGTDDPL**LN** 240

**VVQSESVDLS GLGCGKVLVM VAEKDALVRQ GWGYAAKLEK SGWKGEVEVV ESEGEDHVFH** 300

**LLKPECDNAI EVMHKF**SGFI KGGN 324

**Supplemental Figure S1. Sequence coverage of AtCXE5 and AtCXE12 peptides identified by LC-MS/MS analysis.** Deduced amino acid sequence of AtCXE5 and AtCXE12. Peptides identified by LC-MS/MS analysis are set in bold.

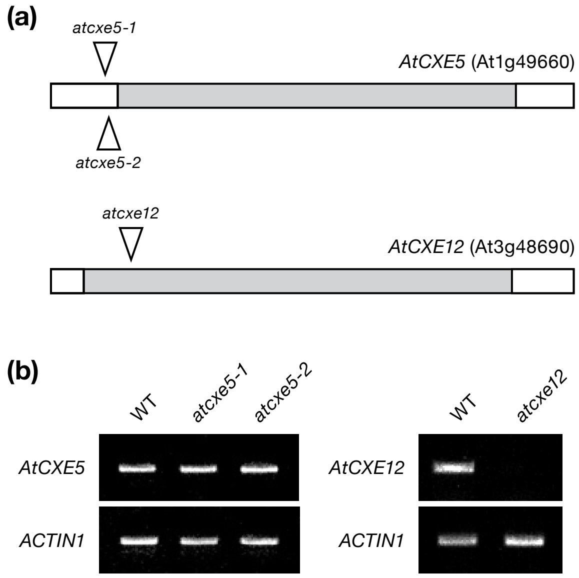


**Supplemental Figure S2. (a)** Structure of the *AtCXE5* and *AtCXE12* loci. Shaded boxes represent untranslated regions; open boxes represent exons. Triangles indicate the location of T-DNA insertions. **(b)** RT-PCR analysis of *AtCXE5* transcripts in leaves from WT and *atcxe5* plants (left panel); RT-PCR analysis of *AtCXE12* transcripts in leaves from WT and *atcxe12* plants (right panel). *ACTIN1* was used as an internal standard.

**Supplemental Table S1. Primers used in this study**

| Name | Sequence (5′–3′) | Purpose |
|---|---|---|
| NdeI-CXE5-F | GACCATATGATGGAATCTGAAATCGCC † | Cloning |
| EcoRI-CXE5-R | GCTATAGAATTCTCAACCAATAATAAACTCGAC | Cloning |
| NdeI-CXE12-F | GACCATATGGATTCCGAGATCGCCGTC | Cloning |
| EcoRI-CXE12-R | GCTATAGAATTCCTAGTTCCCTCCCTTAATAAAC | Cloning |
| SAIL-LB | GCCTTTTCAGAAATGGATAAATAGCCTTGCTTCC | Genotyping |
| SALK-LB | ATTTTGCCGATTTCGGAAC | Genotyping |
| atcxe12-L | CAATGGTACAACCAAACCAAAC | Genotyping |
| atcxe12-R | TCCTGTATCGTTCAAACCAGG | Genotyping |
| atcxe5-1&2-L | CAATGGTACAACCAAACCAAAC | Genotyping |
| atcxe5-1&2-R | TCCTGTATCGTTCAAACCAGG | Genotyping |
| AtCXE5-F378 | AGATGTATGGTCCGCGATTC | RT-PCR |
| AtCXE5-R751 | ATCCCAACCCAGAAAAATCC | RT-PCR |
| AtCXE12-F25 | TGCTCTCCATTGCTCAAAAT | RT-PCR |
| AtCXE12-R498 | GTTTGCTCCTGCACTGTCTC | RT-PCR |
| ACTIN1-F | GAGACAGCCAAAACCAGCTC | RT-PCR |
| ACTIN1-R | TTGATCTTCATGCTGCTTGG | RT-PCR |

† Restriction sites are underlined.

**Supplemental Table S2. List of proteins identified by LC-MS/MS analysis of (*Z*)-3-hexenyl acetate esterase activity purified from Arabidopsis leaves.**

| Accession number | Name | Total intensity | # of spectra | # of unique peptides | Coverage, % |
|---|---|---|---|---|---|
| NP_567103 | Transketolase | 5949487.7 | 151 | 59 | 53.31 |
| NP_187593 | Adenosine kinase 1 | 5365525.9 | 130 | 52 | 59.88 |
| NP_194098 | Polyketide cyclase/dehydrase and lipid transport superfamily protein | 9021641.8 | 154 | 42 | 84.77 |
| NP_190861 | Aldolase superfamily protein | 1627294.6 | 58 | 31 | 65.08 |
| NP_200333 | Trigger factor type chaperone family protein | 1280158.9 | 50 | 30 | 31.63 |
| NP_177875 | PDI-like 1-2 | 1472352.2 | 37 | 23 | 39.96 |
| NP_568203 | ATP synthase alpha/beta family protein | 1258030.6 | 42 | 19 | 30.4 |
| NP_566796 | Glyceraldehyde 3-phosphate dehydrogenase A subunit | 1612348.7 | 51 | 25 | 30.3 |
| NP_190438 | alpha/beta-Hydrolases superfamily protein (CXE12) | 1340105.8 | 45 | 22 | 58.02 |
| NP_001321033 | Ribosomal protein S5/Elongation factor G/III/V family protein | 1033774.6 | 38 | 23 | 19.69 |
| NP_001326777 | Carbonic anhydrase 1 | 1087006.7 | 42 | 17 | 33.98 |
| NP_001189586 | Putative BCR, YbaB family COG0718 | 1029915.4 | 27 | 12 | 48.33 |
| NP_001323480 | Glutathione S-transferase phi 8 | 918498.8 | 41 | 18 | 47.91 |
| NP_195950 | Adenosine kinase 2 | 725480.9 | 29 | 15 | 30.72 |
| NP_192161 | Glutathione S-transferase phi 2 | 770360.1 | 28 | 16 | 51.41 |

| | | | | | |
|---|---|---|---|---|---|
| NP_001189959 | Dehydratase family | 779707.8 | 28 | 15 | 20.79 |
| NP_187884 | Phosphoglycerate kinase 1 | 478440.1 | 20 | 12 | 27.65 |
| NP_175389 | Carboxylesterase 5 (CXE5) | 659779.3 | 22 | 12 | 43.57 |
| NP_001030991 | Ascorbate peroxidase 1 | 596821.2 | 17 | 8 | 33.6 |
| NP_192705 | Oxidoreductase family protein | 744839.5 | 27 | 12 | 26.24 |
| NP_567934 | Pyridoxal phosphate (PLP)-dependent transferases superfamily protein | 361318.7 | 11 | 7 | 20.39 |
| NP_001330235 | Carbonic anhydrase 2 | 393149.3 | 12 | 7 | 23.17 |
| NP_001031721 | Aldolase superfamily protein | 404309.2 | 11 | 7 | 29.33 |
| NP_174996 | Glyceraldehyde-3-phosphate dehydrogenase B subunit | 508541.9 | 22 | 8 | 11.86 |
| NP_194688 | S-adenosyl-L-methionine-dependent methyltransferases superfamily protein | 191745.4 | 6 | 6 | 20.82 |
| NP_174499 | Phospholipid/glycerol acyltransferase family protein | 394318.4 | 16 | 8 | 19.83 |
| NP_197423 | ADP glucose pyrophosphorylase large subunit 1 | 747960.1 | 27 | 13 | 29.31 |
| NP_051066 | ATP synthase CF1 beta subunit | 411085.9 | 15 | 9 | 25.9 |
| NP_191556 | Methylenetetrahydrofolate reductase 1 | 257774.7 | 7 | 5 | 10.98 |
| NP_198035 | Threonyl-tRNA synthetase | 403550.4 | 10 | 5 | 10.86 |
| NP_189966 | Glutathione S-transferase tau 27 | 457196.6 | 17 | 10 | 28.63 |
| NP_181684 | S-formylglutathione hydrolase | 810883 | 19 | 9 | 35.56 |

| | | | | | |
|---|---|---|---|---|---|
| NP_196807 | Selenoprotein O | 754121.8 | 26 | 10 | 8.37 |
| NP_176461 | Stress-inducible protein | 270221.6 | 10 | 6 | 7.88 |
| NP_199588 | Farnesyl diphosphate synthase 1 | 300497.9 | 9 | 4 | 12.5 |
| NP_195780 | Ferretin 1 | 85073.8 | 4 | 3 | 9.8 |
| NP_001030993 | GTP binding Elongation factor Tu family protein | 186865.5 | 8 | 6 | 11.58 |
| NP_566473 | Subtilase family protein | 236228.2 | 8 | 5 | 10.3 |
| NP_176787 | NAD(P)-binding Rossmann-fold superfamily protein | 216487.9 | 9 | 6 | 17.86 |
| NP_001322980 | HISTIDINE TRIAD NUCLEOTIDE-BINDING 2 | 52338.4 | 3 | 2 | 13.61 |
| NP_181187 | Aldolase superfamily protein | 138879.3 | 7 | 5 | 18.99 |
| NP_001031969 | Glutamine synthetase 2 | 548201.2 | 14 | 9 | 22.79 |
| NP_001031599 | Arginase/deacetylase superfamily protein | 187792.6 | 6 | 5 | 19.01 |
| NP_173259 | Calcium-binding EF-hand family protein | 108325.8 | 6 | 5 | 35.88 |
| NP_566316 | Galactose oxidase/kelch repeat superfamily protein | 77981.5 | 5 | 2 | 8.21 |
| NP_194664 | Profilin 2 | 167080.5 | 5 | 2 | 9.92 |
| NP_001077529 | Glyceraldehyde 3-phosphate dehydrogenase A subunit 2 | 257311.8 | 4 | 3 | 5.99 |
| NP_566015 | Calcium-binding EF-hand family protein | 102064.6 | 4 | 3 | 25.35 |

| | | | | | |
|---|---|---|---|---|---|
| NP_194713 | Pyridoxal-5'-phosphate-dependent enzyme family protein | 175863 | 5 | 4 | 10.08 |
| NP_001189932 | Presequence protease 1 | 59284.3 | 3 | 3 | 3.46 |
| NP_190383 | Aldehyde dehydrogenase 2B4 | 420374 | 10 | 4 | 8.18 |
| NP_001324936 | Enolase | 105715.1 | 4 | 2 | 2.77 |
| NP_565178 | Glutathione S-transferase TAU 19 | 181442.6 | 7 | 4 | 12.79 |
| NP_567952 | Cystathionine beta-synthase (CBS) family protein | 162299.2 | 5 | 3 | 17.65 |
| NP_193140 | Selenium-binding protein 2 | 97261.7 | 5 | 3 | 6.37 |
| NP_001078356 | Stromal ascorbate peroxidase | 140099.1 | 4 | 3 | 10.78 |
| NP_199698 | Small nuclear ribonucleoprotein family protein | 40384.4 | 2 | 2 | 18.18 |
| NP_850070 | Nucleotidylyl transferase superfamily protein | 128009.8 | 6 | 3 | 6.13 |
| NP_001318975 | Pyruvate phosphate dikinase, PEP/pyruvate binding domain-containing protein | 256530.4 | 8 | 5 | 5.86 |
| NP_567219 | NFU domain protein 1 | 179212.8 | 4 | 3 | 8.66 |
| NP_568694 | Acetoacetyl-CoA thiolase 2 | 64368.7 | 2 | 2 | 7.69 |
| NP_001184893 | Glutathione S-transferase 6 | 183414.3 | 9 | 6 | 33.17 |
| NP_188056 | Pyridoxal-dependent decarboxylase family protein | 36365.3 | 3 | 2 | 4.13 |

| | | | | | |
|---|---|---|---|---|---|
| NP_200010 | GroES-like zinc-binding alcohol dehydrogenase family protein | 188133.5 | 7 | 4 | 10.99 |
| NP_194791 | Putative BCR, YbaB family COG0718 | 57291.2 | 3 | 2 | 8.33 |
| NP_172153 | Photosystem II subunit P-1 | 46218.8 | 4 | 2 | 8.75 |
| NP_172508 | Glutathione S-transferase family protein | 88041.4 | 5 | 2 | 12.33 |
| NP_565666 | Copper/zinc superoxide dismutase 2 | 93119.2 | 3 | 2 | 12.96 |
| NP_190926 | Diaminopimelate epimerase family protein | 82194.7 | 3 | 3 | 12.15 |
| NP_001321064 | Thylakoidal ascorbate peroxidase | 38189.8 | 3 | 2 | 5.7 |
| NP_196647 | Cystathionine beta-synthase (CBS) family protein | 38279.5 | 3 | 3 | 22.33 |
| NP_001078677 | pfkB-like carbohydrate kinase family protein | 98581.9 | 4 | 2 | 9.06 |
| NP_196867 | Magnesium-chelatase subunit chlH, chloroplast, putative / Mg-protoporphyrin IX chelatase, putative (CHLH) | 158935 | 7 | 5 | 3.84 |
| NP_178480 | Small nuclear ribonucleoprotein family protein | 84660.7 | 3 | 2 | 21.21 |
| NP_190400 | Aldehyde dehydrogenase 10A9 | 40854.7 | 3 | 3 | 9.74 |
| NP_200446 | Pyruvate kinase family protein | 43816.7 | 2 | 2 | 4.02 |
| NP_187678 | Non-intrinsic ABC protein 7 | 98573.1 | 3 | 3 | 14.2 |
| NP_197483 | ARM repeat superfamily protein | 2518420.7 | 20 | 2 | 0.99 |

| | | | | | |
|---|---|---|---|---|---|
| NP_180505 | Glutathione S-transferase tau 6 | 74266.1 | 2 | 2 | 11.21 |
| NP_001318198 | Threonyl-tRNA synthetase, putative / threonine-tRNA ligase | 333314.6 | 6 | 3 | 6 |
| NP_001077966 | Abscisic aldehyde oxidase 3 | 75155.7 | 3 | 3 | 3.83 |
| NP_195870 | Heat shock cognate protein 70-1 | 75644.6 | 3 | 2 | 1.54 |
| NP_001319970 | TIR-NBS-LRR class disease resistance protein | 160869.5 | 3 | 2 | 0.97 |
| NP_001320847 | Ubiquitin carboxyl-terminal hydrolase-related protein | 86703.5 | 3 | 2 | 2.03 |
| NP_174998 | Basic region/leucine zipper motif 60 | 1165346.3 | 10 | 2 | 3.73 |
